# Development of a nucleoside-modified mRNA vaccine against clade 2.3.4.4b H5 highly pathogenic avian influenza virus

**DOI:** 10.1101/2023.04.30.538854

**Authors:** Colleen Furey, Naiqing Ye, Lisa Kercher, Jennifer DeBeauchamp, Jeri Carol Crumpton, Trushar Jeevan, Christopher Patton, John Franks, Mohamad-Gabriel Alameh, Steven H.Y. Fan, Anthony T. Phan, Christopher A. Hunter, Richard J. Webby, Drew Weissman, Scott E. Hensley

## Abstract

Highly pathogenic avian influenza viruses from H5 clade 2.3.4.4b are circulating at unprecedently high levels in wild and domestic birds and have the potential to adapt to humans. We generated an mRNA lipid nanoparticle (LNP) vaccine encoding the hemagglutinin (HA) glycoprotein from a clade 2.3.4.4b H5 isolate. We show that the vaccine is immunogenic in mice and ferrets and prevents morbidity and mortality of ferrets following 2.3.4.4b H5N1 challenge.

## Main Text

Highly pathogenic avian influenza (HPAI) H5 viruses of the A/goose/Guangdong/1996 (Gs/Gd) lineage emerged in southeast Asia in 1996 and have since spread geographically and diversified into several genetically distinct hemagglutinin (HA) clades^1-3^. Long-distance migration of wild birds has enabled rapid transcontinental spread of these HPAI viruses, as evidenced by past H5 outbreaks in 2005-2006, 2014-2015, and 2016-2017^3-5^. Upon re-emerging in 2020, Gs/Gd lineage H5 viruses of clade 2.3.4.4b have circulated at historically high levels in wild and domestic bird populations across Europe, Asia, the Middle East, Africa, and North and South America^2,6-8^. These viruses have persisted with outbreaks continuing uncharacteristically over the summer seasons, wreaking havoc on the poultry industry and resulting in high rates of wild bird mortality^6,9-11^. In comparison to previous H5 outbreaks, a wider range of wild and domestic bird species have been affected by the spread of clade 2.3.4.4b H5 viruses since 2020^12,13^. There have also been occasional human infections and increasing incidences of clade 2.3.4.4b H5 virus spillover into mammals such as red foxes, seals, and minks^14-17^. Some viruses isolated from infected mammals contain genetic mutations associated with mammalian adaptation, highlighting the potential risk an expanded host range can pose^13,15,17-20^.

Our laboratory and others previously demonstrated that mRNA-lipid nanoparticle (LNP) vaccines encoding influenza virus HA induce potent immune responses in mice, rabbits, and ferrets, and clinical trials confirm their safety and immunogenicity in humans^21-24^. We recently developed a multivalent mRNA-LNP vaccine that encodes an HA protein from every influenza virus subtype, including clade 1 H5^24^. This multivalent vaccine protects experimentally infected animals against severe disease and death when challenge strains are antigenically mismatched to the vaccine immunogens^24^; however, the vaccine is not expected to elicit neutralizing antibodies and sterilizing immunity against mismatched influenza virus strains, such as clade 2.3.4.4b H5 viruses. It is therefore important to also develop tailored-made vaccines precisely matched to influenza virus strains with high pandemic potential.

We created a monovalent nucleoside-modified mRNA-LNP vaccine encoding HA from the clade 2.3.4.4b A/Astrakhan/3212/2020 virus. We vaccinated mice with 1 or 10 μg of H5 mRNA-LNP vaccine or 10 μg of a control mRNA-LNP vaccine expressing an irrelevant protein (Ovalbumin), and quantified serum antibody levels and splenic T cell responses after vaccination. Both doses of H5 mRNA-LNP vaccine elicited high levels of HA antibodies that bound (**Figure 1a**) and neutralized (**Figure 1b**) A/Astrakhan/3212/2020. Serum antibody titers declined slightly from 28 to 100 days following vaccination (**Figures 1a-b**). We also tested antibody binding and neutralization of two additional 2.3.4.4b H5 strains, including A/red fox/England/AVP-M1-21-01/2020, which is an H5N8 strain isolated from a red fox, and A/pheasant/New York/22-009066-001/2022 which is an H5N1 virus representative of 2.3.4.4b strains currently circulating in the United States. Relative to the A/Astrakhan/3212/2020 H5 vaccine immunogen, the A/red fox/England/AVP-M1-21-01/2020 HA possesses P152S and S336N substitutions, and the A/pheasant/New York/22-009066-001/2022 HA possesses L120M, V226A, and I526V HA substitutions. The A/Astrakhan-3212/2020-based H5 mRNA-LNP vaccine elicited antibodies that bound (**Figure 1c-d**) and neutralized (**Figure 1e-f**) both A/red fox/England/AVP-M1-21-01/2020 and A/pheasant/New York/22-009066-001/2022. Antibody titers against these variant viruses were ∼3 fold lower compared to A/Astrakhan/3212/2020 titers (**Extended Data Figure 1**). We also examined splenic T cell responses 10 days after vaccination of mice and found that the H5 mRNA-LNP vaccine elicited HA-specific CD8^+^ T cell responses, including CD8^+^ CD44^high^ CD62L^lo^ cells expressing IFNγ (**Figure 1g**) and TNFα (**Figure 1h**).

**Figure 1.**
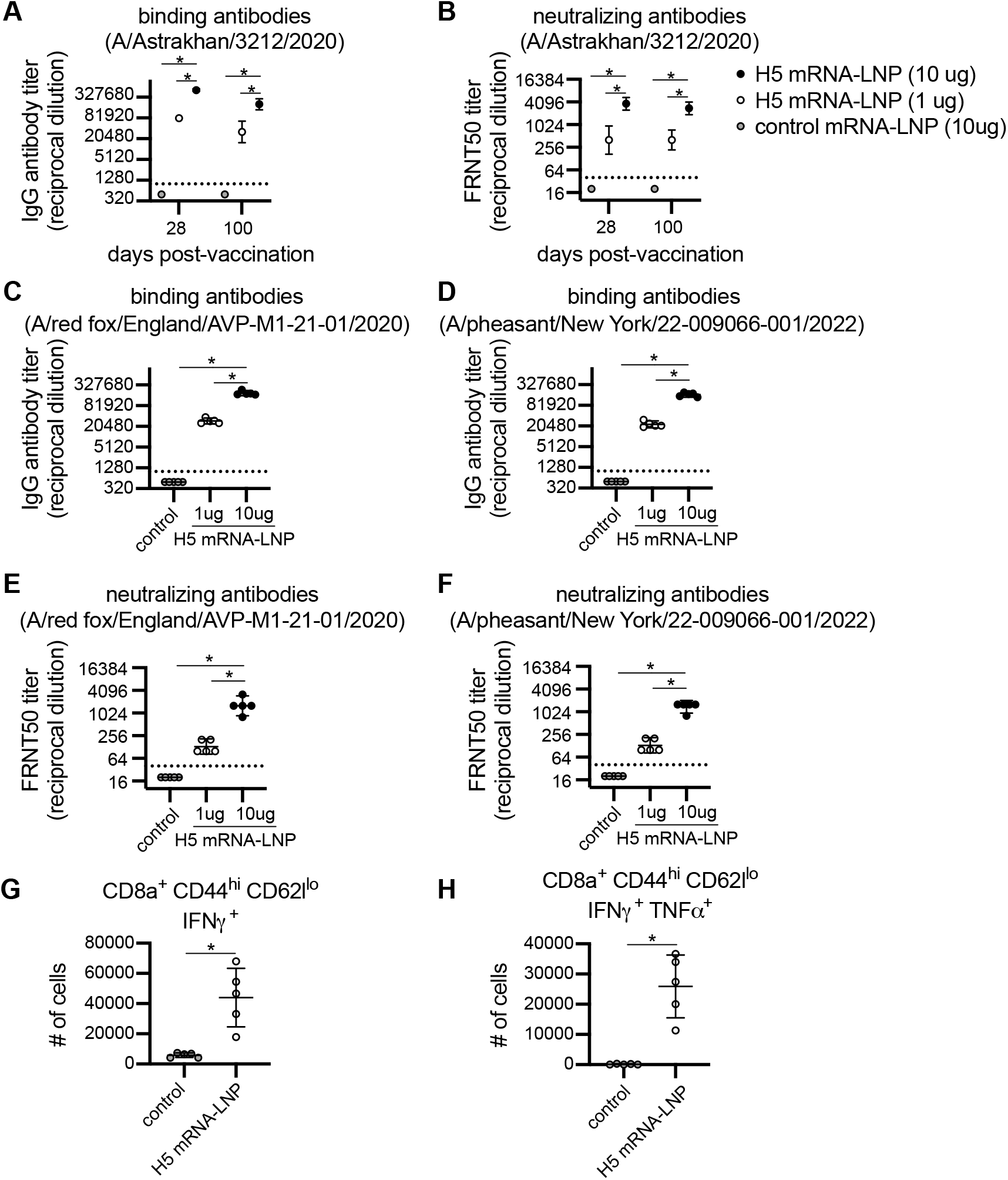
Clade 2.3.4.4b H5 HA mRNA-LNP vaccine induces humoral and cellular immune response in mice. (**A**-**F**) Five mice were included per experimental group. Mice were vaccinated i.m. with 1 or 10 μg A/Astrakhan/3212/2020 HA mRNA-LNP or 10 μg Ovalbumin mRNA-LNP (control mRNA-LNP). (**A**-**B**) Serum samples were collected from mice at 28 and 100 days after vaccination. (**A**) Serum IgG titers reactive to A/Astrakhan/3212/2020 recombinant HA protein were quantified by ELISA. (**B**) A/Astrakhan/3212/2020 neutralizing antibodies were quantified; reciprocal dilutions of serum required to inhibit 50% virus infection are shown. (**C**-**D**) Serum samples collected 28 days after vaccination were tested by ELISA to quantify IgG reactive to A/red fox/England/AVP-M1-21-01/2020 or A/pheasant/New York/22-009066-001/2022 recombinant HA proteins. (**E**-**F**) A/red fox/England/AVP-M1-21-01/2020 and A/pheasant/New York/22-009066-001/2022 neutralizing antibodies were quantified in serum samples collected 28 days after vaccination; reciprocal dilution of serum amounts required to inhibit 50% virus infection are shown. **(G-H)** Five mice per group were vaccinated i.m. with 1 μg A/Astrakhan/3212/2020 HA mRNA-LNP or 1 μg Ovalbumin mRNA-LNP (control mRNA-LNP). Spleens were harvested at 10 days after vaccination and splenocytes were incubated with H5 HA overlapping peptide pools before completing intracellular cytokine staining and flow cytometric analysis of CD8 T cells. Number of IFN-producing CD8^+^ T cells and IFN- and TFN-producing CD8^+^ cells were quantified. All data are shown as geometric means ± 95% confidence intervals. Data in (**A**-**F**) were compared using one-way ANOVA with Tukey’s post hoc test. Values were log-transformed prior to statistical analysis. Data in (**G**-**H**) were compared using an unpaired two-tailed *t* test. Data are representative of 2 independent experiments. *P < 0.05

We vaccinated ferrets using a prime/boost strategy to mimic the dosing schedule initially used for severe acute respiratory syndrome coronavirus 2 (SARS-CoV-2) mRNA vaccination of humans^25,26^. Animals were primed with 60 μg of mRNA-LNP vaccine encoding H5 or an irrelevant protein (Luciferase) and then boosted 28 days later with the same vaccine. Ferrets vaccinated with H5 mRNA-LNP produced high levels of antibodies that bound and neutralized both the A/Astrakhan/3212/2020 (**Figure 2a-b**) and A/pheasant/New York/22-009066-001/2022 (**Figure 2c-d**) viruses. The second dose of H5 mRNA-LNP boosted H5-reactive antibody levels ∼8-fold higher relative to prior to the boost. Antibody titers against A/Astrakhan/3212/2020 and A/pheasant/New York/22-009066-001/2022 were similar (**Extended Data Figure 2**). The vaccinated ferrets were then challenged with A/Bald eagle/North Carolina/ W22-140/2022, an H5N1 strain that is similar to A/pheasant/New York/22-009066-001/2022 and previously shown to be lethal in ferrets^27^. All four H5 mRNA-LNP-vaccinated ferrets survived the H5N1 challenge, while all Luciferase mRNA-LNP-vaccinated animals reached clinical endpoint by 7 days post-challenge (**Figure 2e**). Luciferase mRNA-LNP-vaccinated animals lost more weight (**Figure 2f**), had detectable levels of virus in nasal washes for longer amounts of time (**Figure 2g**), and displayed more clinical signs (**Figure 2h**) relative to H5 mRNA-LNP-vaccinated animals.

**Figure 2.**
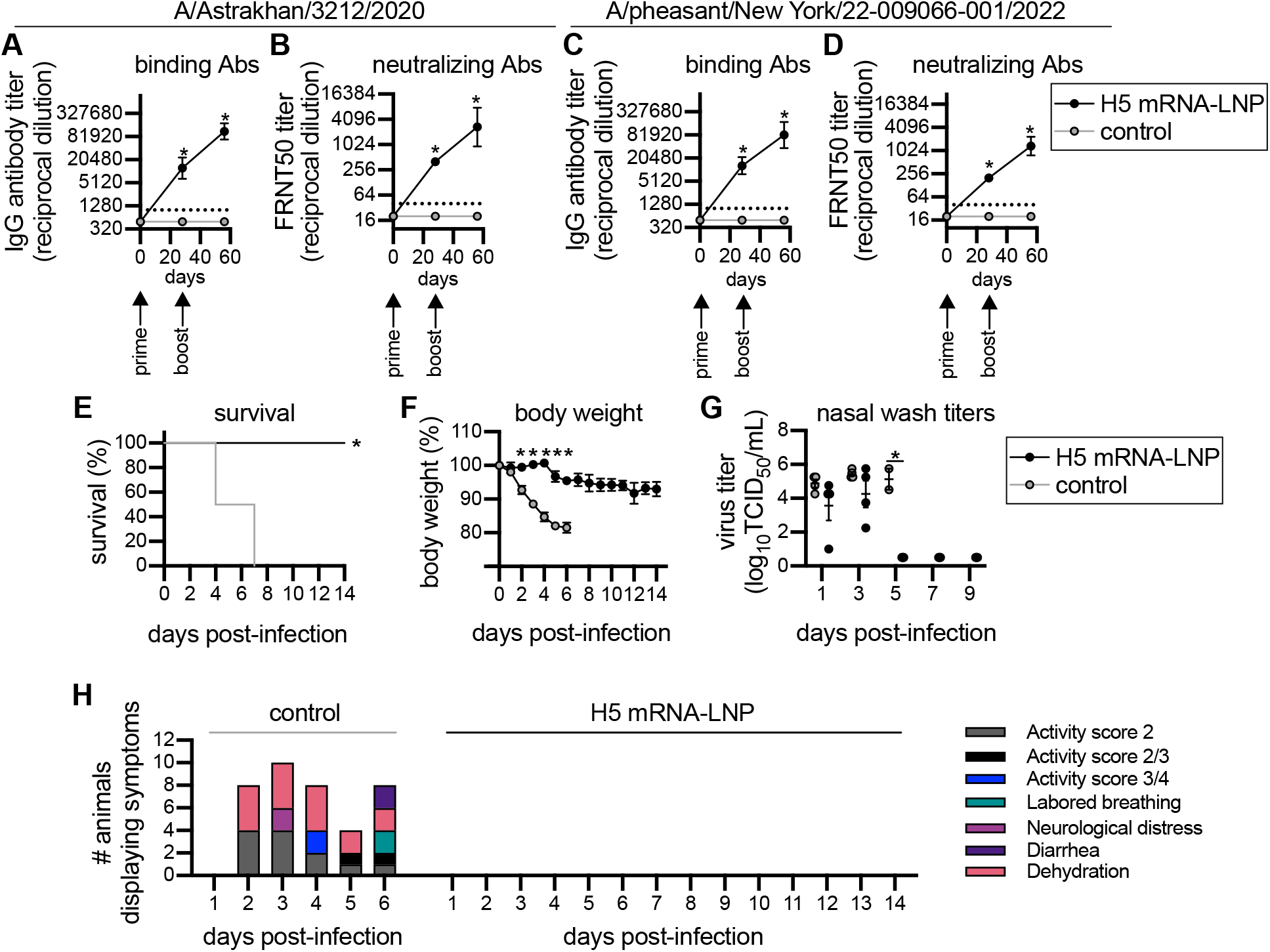
H5 HA mRNA-LNP vaccine protects ferrets from clade 2.3.4.4b H5 virus infection. Four ferrets per group were immunized with 60 μg A/Astrakhan/3212/2020 HA mRNA-LNP or Luciferase mRNA-LNP (control mRNA-LNP), followed by a 60 μg boost 28 days later. Ferrets were challenged with A/Bald eagle/North Carolina/ W22-140/2022 28 days after the second vaccination. Ferret serum samples were collected before vaccination and 28 days after each vaccination. (**A**-**B**) A/Astrakhan/3212/2020 and (**C**-**D**) A/pheasant/New York/22-009066-001/2022 serum IgG binding and neutralizing titers were quantified. Reciprocal dilutions of serum required to inhibit 50% virus infection are shown in panels (**B**) and (**D**). (**E**-**F**) Vaccinated ferrets were monitored for 14 days after i.n. infection with A/Bald eagle/North Carolina/ W22-140/2022. (**E**) Survival, (**F**) body weight, (**G**) virus titers in nasal wash samples, and (**H**) clinical scores are reported after infection. Data in (**A**-**D**) are shown as geometric means ± 95% confidence intervals and values were log-transformed prior to statistical analysis. Data in (**F**-**G**) are shown as means ± SEMs. Data in (**A**-**D, F**-**G**) were analyzed by mixed-model ANOVA with Greenhouse-Geisser correction and Sidak’s multiple comparisons test to compare differences with luciferase (control) mRNA immunization. For animals that died, their weight and viral titers on the day prior to death were carried forward for statistical analyses. Data in (**E**) were analyzed using a log rank test. *P<0.05.

Together, our data demonstrate that our monovalent clade 2.3.4.4b H5 mRNA-LNP vaccine is immunogenic and protective in mice and ferrets. Further studies will evaluate different doses of clade 2.3.4.4b H5 mRNA-LNP vaccine in animal models prior to Phase 1 testing in humans. It will also be important to evaluate the clade 2.3.4.4b H5 mRNA-LNP vaccine in birds and other animals that could potentially serve as an intermediate host to humans, such as swine. Our studies highlight the flexibility of the mRNA vaccine platform which allows for rapid development and precise antigenic matching of vaccine immunogens to emerging influenza virus strains with pandemic potential.

## Funding

This project has been funded in part with Federal funds from the National Institute of Allergy and Infectious Diseases, National Institutes of Health, Department of Health and Human Services, under Contract Nos. 75N93021C00015 (S.E.H.) and 75N93021C00016 (R.J.W.), and grant numbers R01AI08686 (S.E.H.) and R01AI126899 (C.A.H.). Funding was also received from the Commonwealth of Pennsylvania (S.E.H. and C.A.H.) and the Penn Institute for Infectious and Zoonotic Diseases (C.A.H.). S.E.H. holds an Investigators in the Pathogenesis of Infectious Disease Awards from the Burroughs Wellcome Fund.

## Author Contributions

C.F. and S.E.H. designed the experiments, analyzed, and interpreted data and wrote the manuscript. C.F. and N.Y. completed mouse experiments and serological experiments. L.K., J.D., J.C., T.J., C.P., and J.F. completed ferret experiments. A.T.P. completed T cell experiments. M.-G. A., S.H.Y.F., and D.W. designed and produced mRNA-LNP and provided technical advice. C.A.H. supervised T cell experiments and R.J.W. supervised ferret experiments. S.E.H supervised all other activities.

## Competing Interests

S.E.H. and D.W. are co-inventors on patents that describe the use of nucleoside-modified mRNA as a platform to deliver therapeutic proteins and as a vaccine platform. S.H.Y.F. is an employee of Acuitas Therapeutics, a company focused on the development of lipid nanoparticulate nucleic acid delivery systems for therapeutic applications. The authors declare no other competing interests.

