## Extended Data for "Development of a nucleoside-modified mRNA vaccine against clade 2.3.4.4b H5 highly pathogenic avian influenza virus"

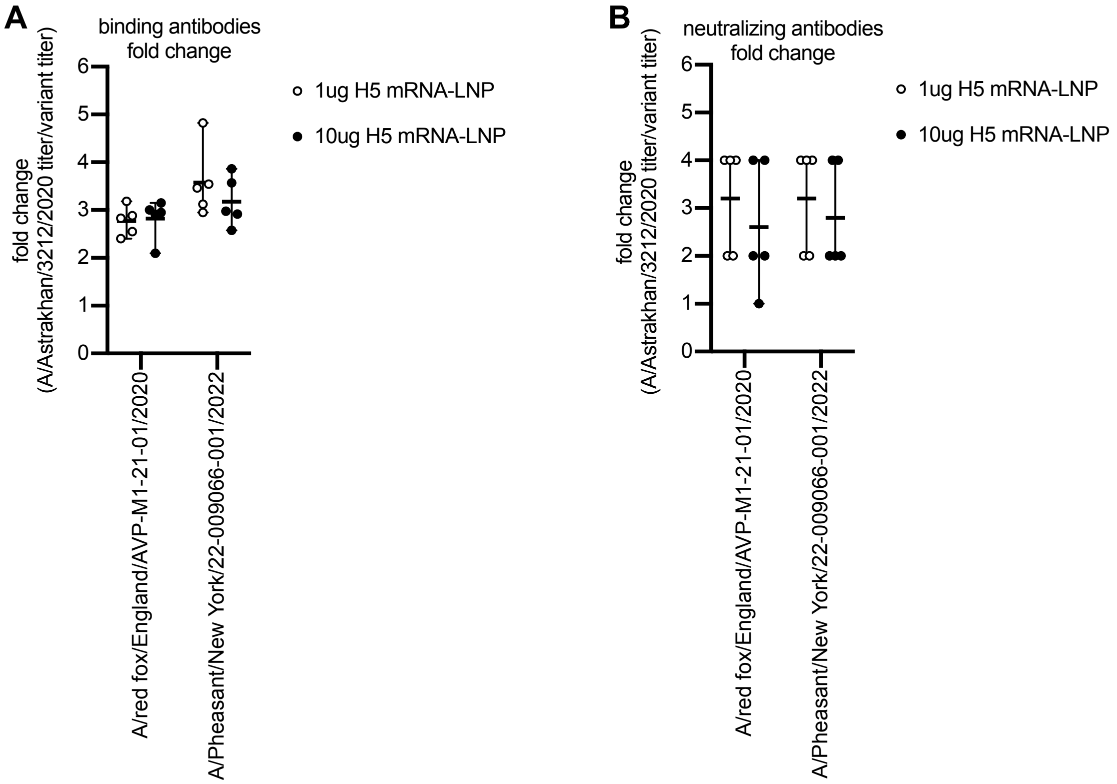


**Extended Data Figure 1. Comparison of murine antibody titers against different Clade 2.3.4.4b H5 antigens.** Serum samples were collected from mice 28 days after vaccination. Antibody binding titers are reported in **Figure 1A**, **C**, and **D** and neutralizing antibody titers are reported in **Figure 1B**, **E**, and **F**. Fold change was determined by dividing antibody titers obtained using A/Astrakhan/3212/2020 antigens with titers obtained using either A/red fox/England/AVP-M1-21-01/2020 or A/pheasant/New York/22-009066-001/2022 antigens. Shown are mean and range.

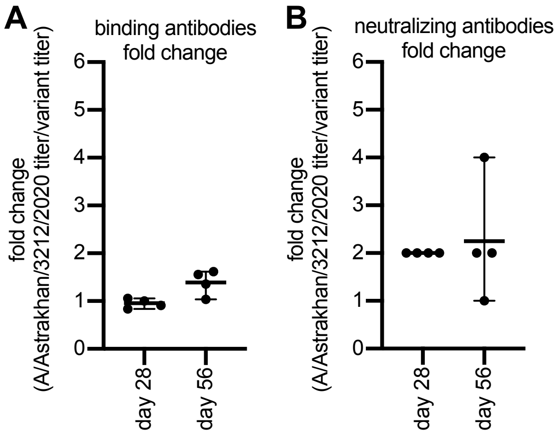


**Extended Data Figure 2. Comparison of ferret antibody titers against different Clade 2.3.4.4b H5 antigens.** Serum samples were collected from ferrets 28 and 56 days after vaccination. Antibody binding titers are reported in **Figure 2A** and **C** and neutralizing antibody titers are reported in **Figure 2B** and **D**. Fold change was determined by dividing antibody titers obtained using A/Astrakhan/3212/2020 antigens with titers obtained using A/pheasant/New York/22-009066-001/2022 antigens. Shown are mean and range.

### Online Methods

#### mRNA-LNP production

The HA sequence of A/Astrakhan/3212/2020 (H5) was codon-optimized, gene synthesized by GenScript, and cloned into an mRNA production vector. mRNAs were then produced as previously described[1] using T7 RNA polymerase (Megascript, Ambion) on linearized plasmids. mRNAs were transcribed to contain 101 nucleotide-long poly(A) tails. Modified nucleoside-containing mRNA was generated using m1Ψ-UTP (TriLink) instead of UTP. *In vitro* transcribed mRNAs were co-transcriptionally capped using the CleanCap (TriLink), and the mRNA purified using cellulose chromatography, as previously described[2]. All mRNAs were analyzed using native agarose gel electrophoresis, and tested to dsRNA and endotoxin content using dot blot and the LAL chromogenic assay respectively before storage at -20°C. Cellulose-purified, nucleoside-modified mRNAs were encapsulated in LNP using a self-assembly process as previously described[3] in which an ethanolic lipid mixture was rapidly mixed with an aqueous solution containing mRNA at pH = 4.0. The LNP used here contain an ionizable cationic lipid, phosphatidylcholine, cholesterol, and polyethylene glycol-lipid. The LNP have a mean hydrodynamic diameter of ~80nm with a polydispersity index of 0.02-0.06 and an encapsulation efficiency of ~95%. The assembled mRNA-LNP were stored at -80°C at a concentration of 1 µg/µL.

#### Mouse experiments

Murine experiments were approved by the Institutional Animal Care and Use Committees of the Wistar Institute and the University of Pennsylvania. Female C57BL/6 mice (Charles River Laboratories) aged 6-8 weeks were immunized intramuscularly with 1-10 µg of mRNA-LNP vaccine encoding either H5 or ovalbumin. For immunizations, mRNA-LNPs were diluted in 50 µl PBS and 25 µl was injected in each hind leg. Blood samples were obtained by submandibular bleeding 28 and 100 days after vaccination and sera were isolated by centrifugation using Z-Gel tubes (Sarsedt). For T cell experiments, spleens were harvested 10 days after vaccination.

**Ferret experiments**

Ferret studies were approved by the St. Jude Children's Research Hospital Institutional Animal Care and Use Committee (IACUC, protocol number 428) in accordance with the guidelines established by the Institute of Laboratory Animal Resources, approved by the Governing Board of the US National Research Council, and carried out by trained personnel working in a United States Department of Agriculture (USDA)-inspected Animal Biosafety Level 3+ animal facility in accordance with all regulations established by the Division of Agricultural Select Agents and Toxins (DASAT) at the USDA Animal and Plant Health Inspection Service (APHIS), as governed by the United States Federal Select Agent Program (FSAP) regulations (7 CFR Part 331, 9 CFR Part 121.3, 42 CFR Part 73.3). This study utilized USDA-classified select agents and A(H5N1) viruses used herein are subject to the guidelines of, and compliance with, requirements discussed in Title 9 (CFR Parts 121 [Possession, Use, and Transfer of Select Agent Toxins] and 122 [Importation and Transportation of Controlled Organisms and Vectors]). Four-to-six-month-old influenza-seronegative male ferrets (Triple F Farms, Sayre, PA, USA) were primed with 60 µg of mRNA-LNP vaccine encoding H5 (n= 4 animals) or an irrelevant protein (Luciferase, n = 4 animals) and then boosted 28 days later with the same vaccine. Blood samples were collected 28 days after each vaccine dose. Animals were lightly anesthetized with isoflurane 28 days after the second vaccination and inoculated intranasally with $10^6$ $EID_{50}$ units of A/Bald Eagle/North Carolina/W22-140/2022 (H5N1) diluted in 1.0 mL of PBS. All animals were monitored daily for clinical signs of infection. Characteristics monitored included body temperature, weight loss, relative inactivity indices, ataxia, respiratory symptoms, stool consistency, and neuropathologic signs. Animals reaching the humane endpoint, according to an IACUC-approved clinical scoring system, were euthanized. Nasal washes were collected from all surviving ferrets at 1, 3, 5, 7, and 9 days post infection (dpi). Ketamine was used to induce sneezing. Viral titers were determined in MDCK cells by $TCID_{50}$ assay.

**Recombinant influenza virus HA proteins**

Recombinant HA (rHA) proteins were generated using pCMV-Sport6 vectors encoding full-length, codon-optimized HA sequences from A/Astrakhan/3212/2020 (EPI ID 1038924), A/pheasant/New York/22-009066-001/2022 (EPI ID 11971502), or A/red fox/England/AVP-M1-21-01/2020 (EPI ID 2081527) with the HA transmembrane domain replaced by the Foldon T4 trimerization domain of T4 fibritin, an Avitag site-specific biotinylation sequence, and a hexahistidine tag as described previously[4]. To produce the recombinant proteins, rHA plasmid and a plasmid encoding neuraminidase (NA) from A/Puerto Rico/8/1934 were co-transfected into 293F suspension cells (Thermo Fisher) using 293Fectin (Thermo Fisher). Supernatants were collected 4 days later for protein purification by Ni-NTA affinity chromatography (Qiagen).

**Enzyme-linked immunosorbent assays (ELISAs)**

ELISAs were performed using 96 well plates (Immulon) coated with 2 μg/mL rHA overnight at 4°C. Blocking and dilution buffer consisted of 1xPBS with 0.1% Tween 20, 0.5% milk, and 3% goat serum. ELISA plates were blocked for 1 hour at room temperature. Heat-inactivated serum samples were serially diluted two-fold in round bottom 96 well plates and were then added to ELISA plates and incubated for 2 hours at room temperature. As a standard control to establish relative serum IgG titers, each ELISA plate included a serial dilution of the CR9114 human monoclonal antibody. Horseradish-peroxidase-conjugated goat anti-mouse IgG (Jackson, 115-035-003; diluted 1:1000), anti-ferret IgG (Abcam, ab112770; diluted 1:2500), or anti-human IgG (Jackson, 109-036-098; diluted 1:5000) were added to ELISA plates and incubated for 1 hour. Plates were developed by adding SureBlue TMB substrate (SeraCare) and quenching the development reaction after 5 minutes using 250 mM hydrochloric acid. Plates were read using a SpectraMax 190 microplate reader (Molecular Devices) at an optical density (OD) of 450 nm.

**In vitro neutralization assays**

Viruses expressing GFP were generated by reverse genetics using the plasmid pHH-PB1flank-GFP that encodes GFP in the open reading frame of the viral polymerase PB1 gene. These viruses can only replicate in cells expressing PB1, as previously described[5].To generate the viruses, a coculture of 293T-CMV-PB1 and MDCK-SIAT1-TMPRSS2-PB1 cells were transfected in serum-free DMEM with 5 bidirectional reverse genetics plasmids encoding PB2, PA, NP, M, and NS from A/Puerto Rico/8/1934 along with pHH-PB1flank-GFP and 2 reverse genetics plasmids encoding the HA and NA segments from clade 2.3.4.4b H5 viruses (A/Astrakhan/3212/2020, A/pheasant/New York/22-009066-001/2022, and A/red_fox/England/AVP-M1-21-01/2020). As an additional biosafety precaution, the multibasic cleavage site KRRKR was edited to a monobasic cleavage site R in the reverse genetics HA plasmids. At 20 hours post-transfection, media was changed to neutralization assay media (NAM) consisting of Medium 199 (Thermo Fisher) with 0.01% heat inactivated fetal bovine serum, 0.3% bovine serum albumin, 100 U of penicillin/ml, 100 µg of streptomycin/ml, 100 µg of calcium chloride/ml, and 25 mM HEPES. Supernatants were collected and clarified by centrifugation at 72 hours post transfection and clarified supernatants were expanded on subconfluent MDCK-SIAT1-TMPRSS2-PB1 cells for an additional 72 hours. Expansion supernatants were collected, clarified, and titrated on MDCK-SIAT1-TMPRSS2-PB1 cells using an EnVision microplate reader at an excitation wavelength of 485 nm and an emission wavelength of 530 nm. Neutralization assays were performed using serum treated with receptor-destroying enzyme (RDE; Denka Seiken) followed by heat-inactivation. Sera were serially diluted two-fold with NAM in flat bottom 96 well plates before an equal volume of virus was added. Serum-virus mixtures were incubated at 37 ºC in 5% $CO_2$ for 1 hour before $2x10^5$ MDCK-SIAT1-TMPRSS2-PB1 cells were added to each well. At 40 hours post-infection, cells were fixed in 4% paraformaldehyde and GFP fluorescence intensities were measured on an EnVision plate reader as described above. 50% neutralization titers are reported as the highest reciprocal serum dilution that decreased GFP levels by 50% or greater in relation to the virus only control wells from each plate, as described previously[6]. Serum samples that were unable to reduce GFP levels by 50% at a 1:40 dilution were given a 50% neutralization titer of 20.

**T cell assays**

A pool of 139 overlapping peptides (15mers, overlapping adjacent peptides by 4 amino acids) spanning the HA sequence of A/Astrakhan/3212/2020 (H5) were prepared (≥70% purity, GenScript) and pooled for *in vitro* restimulation assays. Mice were vaccinated with H5 mRNA-LNP or control mRNA-LNP and spleens were isolated 10 days later. Spleens were processed into single cell suspensions, plated ($5 \times 10^6$ cells/well) and cultured in the presence or absence of the peptide pool (10 µg/mL) for 6 hr at 37°C. Protein Transport Inhibitor Cocktail (Ebioscience: 00-4980-03) was added to cultures after 3 hr for intracellular staining of cytokines and markers of degranulation. Following incubation cultures were harvested and stained for flow cytometry analysis performed on BD FACSymphony A3.

**Quantifying nasal wash viral titers**

Madin–Darby canine kidney (MDCK) cells (ATCC CCL-34) were cultured in Modified Eagle's Medium (MEM) (CellGro) supplemented with 5% fetal bovine serum (FBS) (HyClone), 1 mM L-glutamine, and 1× penicillin/streptomycin/amphotericin B (Gibco) and $NaHCO_3$ (1.5 g/L). Cells were maintained at 37°C in 5% $CO_2$. Infectious viral titers of ferret nasal washes were determined by performing 10-fold dilutions on all nasal wash samples. MDCK monolayers were inoculated with 100 µL of diluted sample and then incubated at 37°C for 72 h. At 72 HPI 50 µL of cell supernatant was mixed with 50 µL of 0.5% chicken red blood cells (CRBC) to measure hemagglutinin agglutination. The endpoint was defined as the highest virus dilution resulting in CRBC agglutination. The 50% tissue culture infectious dose ($TCID_{50}$) titer was determined by the Reed and Muench method[7] and the lower limit of virus detection was 1.0 $\log_{10}$ $TCID_{50}$/mL.
